# Sex- and age-dependent control of dopamine function by LRRK2, and rapid reversal of dopamine release deficits in the LRRK2-G2019S knock-in mouse model of Parkinson’s Disease

**DOI:** 10.64898/2026.10.14.759167

**Authors:** C. Gentle, A. Kamesh, J. Madranges, S. Coady, B. Viera, C. Tifensee-Ribiero, S. McClean, N. Kuhlmann, A. Khayachi, A. Pietrantonio, L. Moulanier, E. Cohen, M. Farrer, M. Parsons, A. Milnerwood

## Abstract

No disease-modifying therapy exists for Parkinson’s disease (PD). At diagnosis (∼65y), ∼70% of nigrostriatal dopamine (DA) axons may have degenerated, so identifying when and how striatal axon function declines is critical. LRRK2 hyperactivity is observed in LRRK2 and VPS35 familial cases, and also idiopathic PD. Thus suggests LRRK2 hyperfunction is a common pathophysiological feature of PD. LRRK2 inhibitors are currently in human trials, yet their effects on brain circuit function are poorly characterized.

By optical imaging of iGluSnFR and dLight we measured striatal glutamate and DA release in WT, LRRK2 G2019S (LKI) and VPS35 D620N (VKI) knock-in mice, across adult ages and sexes, with and without LRRK2 inhibition. In WT mice, ongoing LRRK2 activity regulated DA release in an age- and sex-dependent manner. Acute inhibition reversed basal sex differences in striatal glutamate and DA release in aged mice, and potentiated DA release more in middle-aged females than males. In LKI mice, DA release declined progressively, first in males, then females, after maturation to full adulthood, and remained impaired in very old LKI and VKI mice. Release deficits in aged LKI and VKI mice were resistant to acute LRRK2 inhibition, but 2h inhibition rapidly reversed deficits in middle-aged LKI males and females.

We conclude that LRRK2 activity, and inhibition effects, depend on age, sex, and genotype, with direct implications for the design and timing of current LRRK2 inhibitor clinical trials for PD.

## Introduction

There is currently no disease modifying therapy for Parkinson’s Disease (PD); to provide it we must better understand how neuronal dysfunction in PD occurs. Patients are diagnosed at ∼65y^1^ with mild motor symptoms, by which time ∼70% of nigrostriatal dopamine (DA) axons may have degenerated^2^ and pre-motor symptoms impaired quality of life for years. The best treatment for motor symptoms is DA replacement therapy. While initially effective, this does not help non-motor symptoms or slow progression. Degeneration starts in axons^2^, and may be caused by striatal dysfunction^3^. Early interventions to preserve normal striatal function may prevent or slow degeneration, whereas interventions after onset of degeneration may need to be different. Our goal is to identify when, and how striatal axon function declines, and devise strategies to prevent it.

Approximately 1/4 of patients have a family history of PD^1^, and genetics underly ∼30% of population risk^4^. Mendelian discoveries, supplemented by evidence of environmental and other risk factors, are starting to reveal the molecular evolution of PD and may help explain when, why, and how the disease occurs^5^. Knowing how gene variants alter axon function before degeneration might show us how to prevent it. The most common PD causal mutations are in the kinase LRRK2^6,7^; they all increase LRRK2 activity and hyperphosphorylate its substrates^8^. A rarer, but highly penetrant, PD mutation in VPS35 also causes hyperphosphorylation of LRRK2 substrates^9,10^, and LRRK2 substrate hyperphosphorylation also occurs in idiopathic PD patient brain (with no known mutations)^11^. This suggests LRRK2 hyperactivity is a common pathophysiological stress in PD onset.

Accordingly, LRRK2 inhibitors have entered human trials; however, literature on the effects of LRRK2 inhibition on brain circuit function, and mutant dysfunction, is lacking. Here we demonstrate that ongoing LRRK2 activity regulates DA release in an age- and sex dependent manner; acute inhibition of LRRK2 consistently reversed basal sex differences in striatal glutamate & DA release properties in aged WT mice, and had a greater potentiating effect on DA release in middle aged females than males.

Since our first reports of LRRK2 G2019S & VPS35 D620N PD knock-in mice (LKI & VKI)^12–14^ we and others showed both models exhibit some PD hallmarks e.g., synuclein pathology or accumulation, aberrant tau phosphorylation, nigral cell loss, and LRRK2 hyperphosphorylation^13–18^. In young-adult LKI & VKI^14,19^ we found evidence of increased DA release, which could be an early pathophysiological toxic stress. In support of this notion, similar changes (increased DA turnover) have been found in asymptomatic LRRK2 PD mutation carriers, prior to onset of clinical PD^20^.

We recently found increased DA release in young-adult VKI mice was reduced by chronic or acute LRRK2 inhibition^18,21^; suggesting LRRK2 inhibitors normalize increased release in young-adult VKI mice. This would potentially be useful for early PD, if it removes a toxic stress. Reducing DA release in scenarios where it is already impaired would likely be deleterious.

Here we show DA release is progressively reduced in LKI mice, first in males, then females. Impairments emerge after maturation to full adult ages, and remain impaired in very old LKI & VKI mice, at ages analogous^22^ to when people present with PD^1^. We found release deficits in aged LKI & VKI mice were resistant to acute LRRK2 inhibition. However, LRRK2 inhibition rapidly (2h) reversed impaired DA release in middle-aged male and female LKI mice.

Knowing what LRRK2 does in adult, and very old WT, LKI, & VKI mice is of basic science interest, and the age- and sex-effects we uncovered have immediate implications for LRRK2 inhibitor clinical trials for PD.

## Materials & Methods

### Animals

LRRK2 G2019S (LKI) & VPS35 D620N (VKI) knock-in mice were generated as previously described (Beccano-Kelly et al., 2014; Yue et al., 2015)^10,18^ and maintained on a C57Bl6/J wild-type (WT) background. Mice were housed and bred in accordance with the Canadian Council on Animal Care regulations (Animal Use Protocol 2017-7888B). All procedures were approved by and governed in accordance with the Neuro Centre of Neurological Disease Models (Animal Use Protocol MNI-7888). To experimentally model asymptomatic, prodromal, end-stage of PD, 3-, 6-, & 25-month-old male and female WT, heterozygous LKI & VKI mice were used for all measures of glutamate and DA transmission. Cross-species lifespan analysis approximates 3-month mice to be young adults (∼20yrs), 6-month-old mice to middle age (30-40y), and 25-month mice analogous to ∼65y^22^, when people typically present with idiopathic & LRRK2 PD^1^..

### Genotyping

Genotyping was conducted on all mice by ear tissue at weaning and confirmed post-mortem. Briefly, samples were digested in 100 µL 10% Chelex (Bio-Rad 142–1253; 20 min 95 °C, 2x vortex) and centrifuged (2 minutes 12,000 RPM) before DNA-containing supernatant (2.5 µL) was added to PCR master mix (17.5 µL, Qiagen 201203: taq polymerase, DNAse and RNAse-free water, 10X buffer, 10 mM dNTPs, and custom DNA oligo primers as previously ^10,12,18,19,23,24^. DNA was amplified by PCR and combined with fluorescent DNA intercalating dye (ZmTech LB-001G) prior to electrophoresis. 10 µL of PCR product was run in 1.5% agarose gel and visualized (BioRad UV gel imager) for the presence of 1 or 2 bands to determine WT vs LKI or VKI genotype, respectively.

### Surgery

At ∼2.5, 5.5, & 24.5 months of age, mice received stereotaxic injection of viral constructs to express dLight1.3b, a genetically encoded fluorescent DA reporter (AAV5-CAG-dLight1.3b, Addgene 125560-AAV5, 1.5 × 1013 GC/mL) and / or iGluSnFR, a genetically encoded glutamate sensor (AAV9.hSyn.iGluSnFr.WPRE.SV40 (Addgene 98929-AAV9, 2.8 × 1013 GC/mL) injected (300 nL, 1nL/sec) into the dorsolateral striatum (coordinates: +1.0 mm anterior, +/-2.0 mm lateral, -3.2 mm ventral), as previously^18^.

### Preparation of acute brain slices

300 µm acute coronal brain slices were prepared from restrained mice which were swiftly decapitated. Brains were quickly transferred to ice-cold recovery solution (1 min; in mM: 93 NMDG, 93 HCl, 2.5 KCl, 1.2 NaH2PO4, 30 NaHCO3, 20 HEPES, 25 glucose, 5 sodium ascorbate, 3 sodium pyruvate, 10 MgSO4·7H2O, 0.5 CaCl2·2H2O, pH 7.3-7.4, 290-310 mOsm; carbogen-infused). Following this, extra-striatal rostral and caudal brain regions were removed by coronal sectioning, and the remaining block was mounted onto a vibrating-blade microtome platform with sodium acrylate (Leica Micro-systems VT 1200S). After hemisectioning at the midline, coronal sections were cut, and slices were then transferred to warm recovery solution (35°C,15 min). A final transfer of slices into holding chambers containing room-temperature artificial cerebrospinal fluid (aCSF; in mM: 125 NaCl, 3 KCl, 25 NaHCO3, 1.25 NaH2PO4, 2 MgCl2, 2 CaCl2, 10 glucose, pH 7.2-7.4, 300–310 mOsm, 22–25 °C, carbogen-infused) was followed by ≥45min settling period, or 2h for drug preincubation, as previously^18^.

### Imaging

dLight & iGluSnFR video recordings were obtained using a 4X objective focused on the DLS. Slices were wide-field illuminated with blue light (473 nm, CoolLED pE-340fura), with the shutter, EM-CCD camera (Andor iXon Ultra 897), and stimulus isolator (WPI A365) triggered by Clampex & Digidata 1550B). The stimulus isolator was connected to a monopolar tungsten stimulating electrode (A-M Systems 574000) lowered into dorsolateral striatum 50–100 µM beneath the surface of the tissue. Recordings were captured using Andor Solis software using 4 × 4 binning at 205 Hz. For these experiments, spontaneous dLight transients were excluded from analysis of responses to electrical stimulation. 2 or more pulses (at 4s or 100ms) inter-pulse intervals were delivered at 50-400uA, every 2min to generate stimulus response curves. To account for changes in background fluorescence and photobleaching, no-stimulation and stimulation trials were alternated at 1-minute intervals, and repeated every 2min, until 6 no-stimulation trails were recorded, as previously^18^.

### LRRK2 inhibition

LRRK2 kinase activity was inhibited by the selective LRRK2 inhibitor MLi-2 (Tocris 5756), as previously ^10,18^ in the cyclodextrin diluent Captisol® (Ligand RC-0C7-100). MLi-2 stock solution was prepared by bath sonicating 1 mg MLi-2 in 2 mL 45% Captisol®-PBS for approximately 3 h at 4°C, followed by continuous agitation on a rotary mixer overnight until fully dissolved. Asides from IP injection of aged mice, brain slices were paired by hemisphere, one maintained in aCSF containing 45% Captisol® (vehicle condition), or aCSF containing 45% Captisol® + MLi-2 (500nM; MLi-2 condition). All slices were initially held in vehicle aCSF, after which treatment slices were transferred to vehicle or MLi-2 bath for 2h. Following treatment, slices were recorded while continuously perfused with their respective vehicle or MLi-2-containing aCSF.

### Protein quantification immunoassays

Protein expression levels were analyzed in striatal brain slices or whole brain as indicated. Samples were flash frozen in liquid nitrogen and stored at -80°C after recording DA release. Protein lysates were prepared in ice-cold tris buffered saline (TBS; 0.1% Tween 20) containing a cocktail of protease inhibitor (Roche 11697498001) and phosphatase inhibitor (Sigma 4906845001). Samples were homogenized by probe sonication for 15s at 70% intensity, rotated at 4°C for 30 min, and then centrifuged for 10 min at 7000 RMP.

For fluorescence western blotting, 30 μg of protein was prepared in 4× NuPAGE LDS sample buffer (Invitrogen, NP0008) containing 2.5% β-mercaptoethanol and denatured at 95°C for 5 min prior to loading onto Bolt 4–12% Bis–Tris Plus gels (Invitrogen, NW04120BOX). Proteins were separated by electrophoresis at 70 V for 30 min, followed by 110 V for 1 h, and 2.5 h for larger proteins, including LRRK2. Proteins were then transferred onto Immobilon-FL PVDF membranes (Millipore, IPFL00010). Membranes were blocked with TBST (TBS, 0.1% Tween-20, 5% BSA) for 1 h at room temperature and incubated with primary antibodies (Rab12, Proteintech 18843-1-AP, 1:1000; pRab12, Cell Signaling 45036, 1:1000; pLRRK2, Abcam ab203181, 1:1000; GAPDH, ThermoFisher MA5-15738, 1:2000) overnight at 4 °C. TBS-based buffers were used throughout to minimize potential interference with phospho-specific antibodies. Following three 10 min washes in TBST, membranes were incubated with fluorescent secondary antibodies (LI-COR) for 1 h at room temperature and imaged using a LI-COR Odyssey Infrared Imaging System (LI-COR).

When necessary, phosphorylated or low-abundance proteins were probed first, after which membranes were stripped using LI-COR NewBlot IR Stripping Buffer (LI-COR, 928-40028) in 30 min intervals until the initial signal was no longer detectable. Membranes were subsequently reprobed for non-phosphorylated or higher-abundance proteins of similar molecular weights. Images were background-subtracted, and band intensities were quantified using Image Lab software. Protein signals were normalized to a housekeeping protein (GAPDH, β-tubulin, or β-actin) quantified from the same gel.

Lysates were also quantified with a Protein-Simple Wes automated capillary-based size sorting system as previously ^10,12^. Briefly, lysates were diluted to a final concentration of 0.5 µg/µL in 0.1X sample buffer (Bio-techne 042-195) and combined with 5X fluorescent master mix (Bio-techne PS-FL01-8) in a 4:1 ratio. Primary antibodies (as above), Wes antibody Diluent II (Bio-techne 042-203), horseradish peroxidase (HRP)-conjugated secondary antibodies (Bio-techne 042-205 & 042-206), streptavidin-HRP (Bio-techne 042-414), a 1:1 Luminol-Peroxide mix (Bio-techne 043-311 & 043-379) and wash buffer (Bio-techne 042-202) were then loaded. The assay plate was centrifuged at 1,000 x g (∼2,500 rpm) for 5 minutes and loaded into the Wes instrument with a 25-capillary cartridge (Bio-techne PS-CC01). Protein separation by molecular weight, UV-light cross-linking immobilization to the capillary wall, immunoprobing, and chemiluminescent detection were performed automatically according to the manufacturer’s protocol. Data were analyzed using Compass for Simple Western software, with target protein signals normalized to GAPDH.

### Analysis & Statistics

Fluorescence imaging data were analyzed using FIJI and/or a semi-automated custom Python-based pipeline, EMMA (Eli’s Marvelous Mouse Analysis. Code available on request). TIFF image stacks were processed using Python libraries including NumPy, Pandas, SciPy, tifffile, and Matplotlib. Circular regions of interest (ROIs) were manually selected for each slice, and mean fluorescence intensity across the z-axis was extracted to generate fluorescence profiles. Baseline and peak fluorescence values were quantified from predefined analysis windows, and release transients quantified as dF/F (%). Paired-pulse ratios (PPRs) were calculated as the ratio of P2 to P1 dF/F responses. Exponential decay kinetics (tau) were determined by fitting transients with decay curves, using nonlinear least-squares exponential regression. Exclusion criteria were determined using a data-driven approach based on distribution-based thresholding methods. Data was excluded based on the following parameters: (1) baseline drift exceeded a predefined threshold, determined using the interquartile range method (Q3 + 1.5xIQR); (2) abnormal baseline fluorescence values, corresponding to values outside the 2.5-97.5 percentile range; (3) low evoked responses (max P1 dF/F% <10% dLight, <2% iGluSnFR), below the reliably detectable signal range. All data curation and statistics were conducted in GraphPad Prism11. Non-parametric data were analyzed using Kruskal-Wallis testing. Parametric data were analyzed using 1-way or 2-way ANOVA, with Holm-Šídák or FLSD multiple comparisons tests used for post-hoc analyses when ANOVA reached significance (p < 0.05). Pairwise comparisons were performed using Fisher’s LSD test where indicated. Asterisks represent significance of pair-wise comparisons *p < 0.05, **p < 0.005, ***p < 0.0005, ****p <0.0001. Trends were defined as 0.15 > p > 0.05. All statistical analyses are described in figures & legends. Data are presented as n=observations from (n) animals (e.g., WT=6(3) is 6 slice observations from 3 WT animals).

## Results

### Dopamine release deficits in LKI mice emerge earlier in males

We recently reported that VPS35 D620N (VKI) mice exhibit LRRK2 hyperphosphorylation, and increased DA release (assayed by dLight) at 6, but not 3 months. Increased DA release was rapidly reversed by LRRK2 inhibition ^18^. Here we conducted similar investigation of LRRK2-G2019S (LKI), employing ultrafast imaging of the genetically encoded DA sensor dLight in acute dorsolateral striatal (DLS) of coronal slices from 3- and 6-month-old WT & LKI mice (Figure 1A). DA release was evoked by paired electrical stimuli delivered with a 4 s inter-pulse interval (IPI), and increasing stimulation intensities (Figure 1Aiii). At 3 months, dLight response transient peak amplitudes did not significantly differ between WT and LKI mice (Figure 1Bi). However, by 6 months, LKI mice exhibited significantly reduced evoked DA release, relative to age-matched WT controls (Figure 1Bii). When segregated by sex, no significant genotype differences were observed in either males or females at 3 months (Figure 1Biii,v). In contrast, at 6 months, DA release was significantly reduced in LKI males relative to WT males, while no significant genotype differences were observed in females (Figure 1Biv,vi). These findings indicate that deficits in evoked DA release emerge between 3 and 6 months of age in LKI mice, with males exhibiting greater susceptibility to early dopaminergic dysfunction.

**Figure 1.**
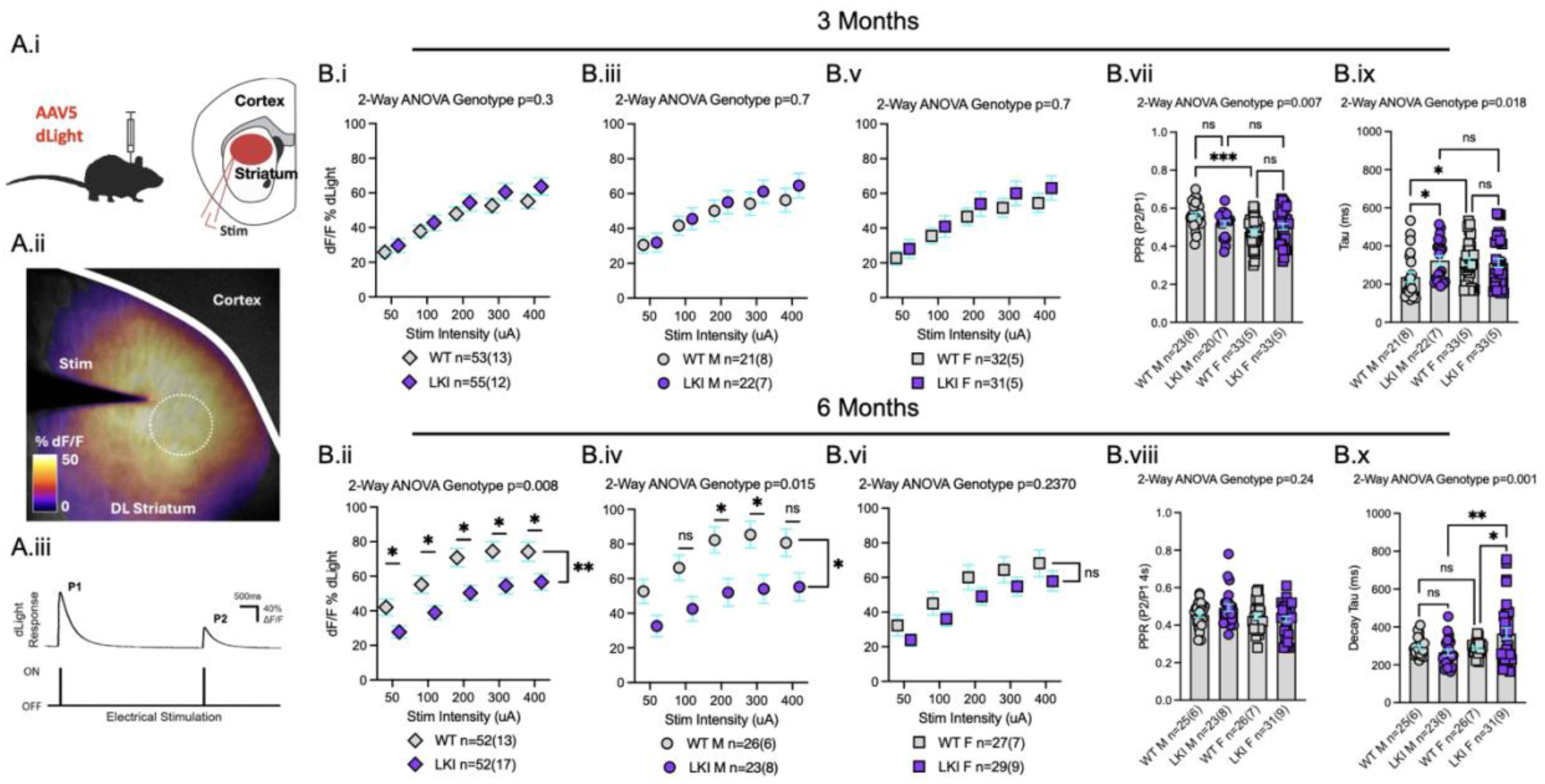
Dopamine release deficits develop earlier in male LKI. **A.i)** Schematic depicting stereotaxic injection of AAV5-dLight to dorsolateral striatum (DLS) 3–6 weeks prior to acute slice preparation, and cartoon depicting dLight expression (red) in coronal slices containing cortex and striatum, with local stimulation delivered to the DLS. **ii)** Example still from CMOS ultrafast imaging of dLight fluorescence evoked by local electrical stimulation, overlaid with colour lookup table showing %dF/F from baseline. Solid white line denotes the corpus callosum separating cortex and striatum, dashed circle indicates typical 40-pixel ROI used for quantification, centered on hotspot. **iii)** Representative dLight transient responses evoked by 2 pulses (P1 & P2) delivered with 4s inter-pulse interval (IPI). **B.i&ii)** At 3 months of age (young adult) there was no significant difference between WT & LKI mice in P1 response peaks to increasing stimulation intensity (i), whereas DA release is significantly reduced in slices from 6 month LKI relative to WT littermates of the same age(ii). **iii-vi)** Segregation by sex showed no change to peak release in male (iii) or female (v) mice at 3 months. In 6-month-old animals. a large reduction of DA release is observed in males (iv), but not females (vi). **vii&viii)** Paired-pulse ratios (PPR; P2/P1) are lower in female WT mice, relative to males at 3 months; this sex difference is not observed in LKI (vii) and is absent in WT & LKI at 6 months (viii). WT PPRs reduce in both sexes between 3 (vii) and 6 months (viii), together suggesting female DA release properties may mature more rapidly than males (lower than males at 3 months). **ix&x)** P1 transient decay tau is higher (slower) in female WT mice, relative to males at 3 months; this sex difference is not observed in LKI (ix) and is absent in WT at 6 months (x). LKI decay tau is higher in 3-month male LKI relative to WT males (when release deficits are yet to occur) and female decay tau is not different from female WT (ix). At 6 months, male LKI decay tau is no longer different from WT, and female decay tau is slower than WT females & LKI males (x). Together, data suggest slower decay tau precedes reductions in DA release peaks.

To investigate the mechanisms that regulate presynaptic DA release, we also examined paired-pulse ratios (PPRs; P2/P1) at both ages. At a 4s IPI, DA release typically displays paired-pulse depression, widely held to be indicative of re-release capacity and DA availability, and the inhibitory action of presynaptic D2 auto-receptors on nigral axons^25^. At 3 months, WT females exhibited significantly lower PPRs than WT males; this sex difference was absent in LKI mice (Figure 1Bvii). By 6 months, no significant sex differences in PPR were observed in either genotype (Figure 1Bviii). PPRs were lower in male and female WT mice at 6 months, relative to 3 months, suggesting age-dependent reduction / maturation of the presynaptic mechanisms regulating DA release.

DA response kinetics (tau) are influenced by stochastic release and to a large extent DA reuptake / clearance by the dopamine transporter (DAT)^25^. We quantified DA decay constants and found WT females exhibited significantly slower decay kinetics than WT males at 3 months; again, this basal sex difference was absent in LKI (Figure 1Bix). Notably, decay tau was significantly increased in 3-month-old LKI males, relative to WT males, despite no detectable differences in peak DA release at this age. By 6 months, when release amplitude was reduced in male LKI, decay tau no longer differed between LKI and WT males. At this age, LKI females exhibited significantly slower decay kinetics than both WT females and LKI males (Figure 1Bx), and a trend toward reduced release. The results show progressive age- and sex-dependent effects on striatal DA neurotransmission in LKI mice, with slower DA decay kinetics preceding the emergence of DA release amplitude deficits.

### LRRK2 hyperphosphorylation is sustained in LKI mice

To assess LRRK2 kinase activity, we performed western blot analysis of phosphorylated LRRK2 (pLRRK2; S1292) and its well characterized substrate Rab12 (pRab12; S106)^8^. In striatal slices from 6-month-old WT and LKI mice (Figure 2A,B) we found significantly increased pLRRK2:LRRK2 in male and female LKI mice, relative to WT (Figure 2Aii). Similarly, pRab12:Rab12 was significantly increased in both male and female LKI mice, indicating elevated LRRK2 activity & substrate phosphorylation at 6 months (Figure 2Bii).

**Figure 2.**
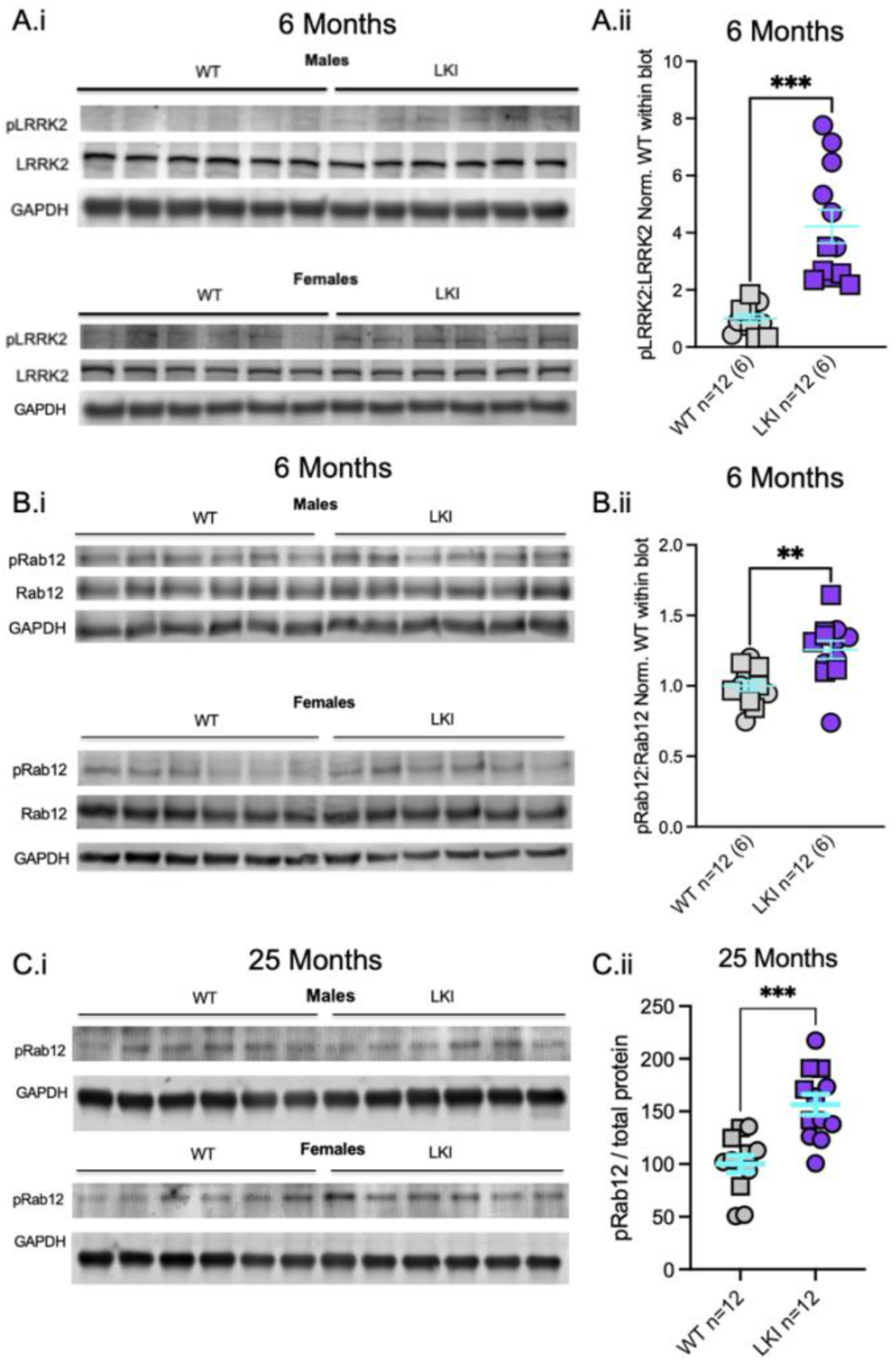
LRRK2-dependent hyperphosphorylation in LKI striatum. **Ai)** Striatal brain slices from 6-month mice collected after recording were subject to western blot of phosphorylated LRRK2 (S1292; pLRRK2), total LRRK2, & GAPDH. **ii)** Quantification of pLRRK2:LRRK2 in male (circles) and female (square) tissue, normalized to WT within blot. pLRRK2 is elevated in male & female LKI striatum. **Bi)** Western blot of LRRK2-dependent Rab12 phosphorylation (S106; pRab12), total Rab12, & GAPDH. **ii)** Quantification of pRab12:Rab12 in male (circles) and female (square) tissue, normalized to WT within blot. pRab12 is elevated in male & female LKI striatum. **Ci)** Western blot of pRab & GAPDH in brain tissue of 25-month-old WT & LKI mice. **ii)** Quantification of pRab12:total protein (stain not shown) in male (circles) and female (square) tissue, normalized to WT within blot. pRab12 is elevated in male & female LKI brain.

To determine if elevated LRRK2 activity persists with ageing, we examined pRab12 in brain tissue from 25-month-old WT and LKI mice (Figure 2C). pRab12 (normalized to total protein) revealed significantly elevated phosphorylation in both male and female LKI mice, relative to WT controls (Figure 2Cii). The data indicate elevated LRRK2 activity across the lifespan of LKI mice.

### Dopamine and glutamate neurotransmission are impaired in aged LKI and VKI males and females, and are resistant to LRRK2 kinase inhibition

To investigate striatal neurotransmission and LRRK2 kinase inhibition in aged mice, we conducted imaging of iGluSnFR and dLight to quantify examined evoked glutamate and DA release in 25-month-old WT, LKI, and VKI mice (Figure 3A). 2 h prior to acute slice preparation, animals received an intraperitoneal injection of vehicle (Captisol®) or MLi-2 (5 mg/kg), with slices subsequently maintained in vehicle or 500nM MLi-2 throughout recording.

**Figure 3.**
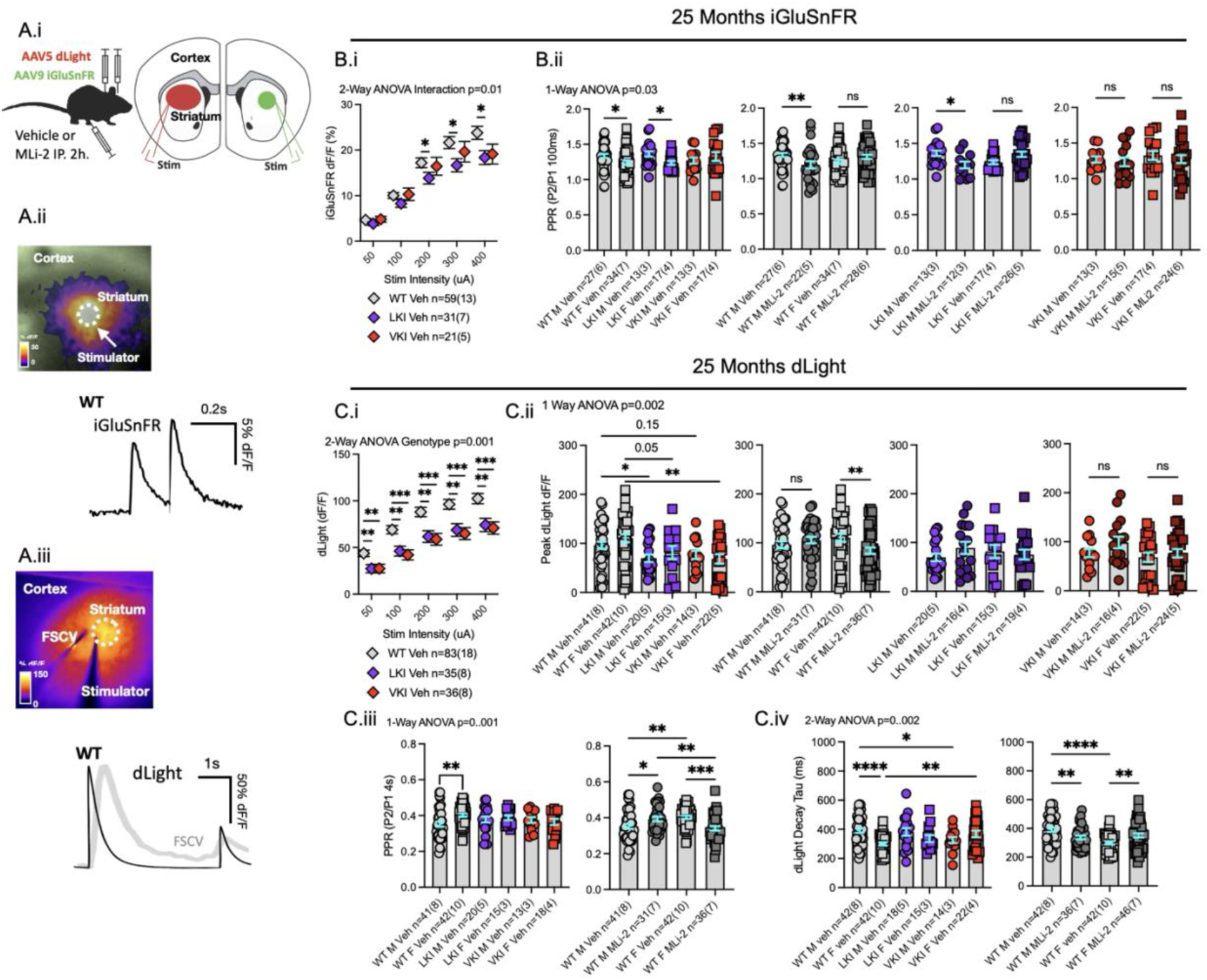
Sex-dependent effects of LRRK2 inhibition on glutamate & dopamine release in 25-month WT mice, and no rescue of release deficits in LKI & VKI. **A.i)** Schematic depicting stereotaxic injection of unilateral AAV5-dLight & AAV9-iGluSnFR to each dorsolateral striata (DLS) 3–6 weeks prior to acute slice preparation, and cartoon depicting dLight (red) & oGluSnFR expression in coronal slices of cortex and striatum, with local stimulation delivered to the DLS. Animals received intraperitoneal (IP) injection of vehicle (Captisol®) or MLi-2 (5mg/kg) 2h prior to brain slicing, and slices were maintained in vehicle or 500nM MLi-2. **ii)** Example still from CMOS ultrafast imaging of iGluSnFR fluorescence transient evoked by local electrical stimulation, overlaid with colour lookup table showing %dF/F from baseline. Dashed circle indicates typical 40-pixel ROI used for quantification, centered on hotspot with representative iGluSnFR transient responses evoked by 2 pulses delivered with 100ms inter-pulse interval (IPI). **iii)** Example still of dLight fluorescence evoked by local electrical stimulation, overlaid with colour lookup table showing %dF/F from baseline. Dashed circle indicates ROI used for quantification, centered on hotspot alongside representative dLight transient response evoked by 2 pulses (P1&P2; 4s IPI) with representative fast-scan cyclic voltammetry trace for comparison. **B.i)** At 25 months (aged) there was a significant interaction between genotype and stimulation intensity, due to reduced iGluSnFR peak amplitude in response to increasing stimulation intensity in LKI & VKI mice, relative to WT. Multiple comparison post-testing revealed significant reductions at higher stimulus intensities in LKI mice. Sex differences in peak amplitudes were not apparent (not shown). **ii)** Glutamate PPRs (100ms IPI) were not significantly different between WT, LKI, & VKI mice, but were significantly reduced in WT & LKI females, relative to males, indicating higher probability of release in females. This sex effect was absent in VKI striata. Statistical analysis included all comparisons but are separated for clarity. LRRK2 inhibition significantly reduced glutamate PPRs in WT males, but not females, reversing the sex-differences in glutamate PPRs. This pattern was maintained in LKI, but not VKI mice. **C.i)** There was a highly significant main genotype effect on DA release in LKI & VKI mice, relative to WT, at all stimulation intensities. Sex differences were not apparent (not shown). **ii)** There was a highly significant ANOVA effect of group (sex, genotype & treatment) on DA release maxima. Statistical analysis included all comparisons but are separated for clarity. There were no differences to DA release maxima between vehicle treated male & female mice of any genotype. Release was significantly reduced in LKI males & VKI females, relative to WTs, and strongly trending to significant reductions in LKI females and VKI males. LRRK2 inhibition significantly reduced DA response maxima in WT females, but not males. This pattern was not maintained in LKI or VKI mice. **iii)** There was a highly significant ANOVA effect of group (sex, genotype & treatment) on DA PPRs. Statistical analysis included all comparisons, but data are separated for clarity. DA PPRs (4s IPI) were significantly higher in female than male WT mice, but there were no differences from, and no sex differences within LKI & VKI mice. LRRK2 inhibition had no effect on LKI or VKI PPRs (not shown) but significantly increased and decreased male & female WT PPRs, respectively, reversing base sex differences. **iv)** There was a highly significant ANOVA effect of group (sex, genotype & treatment) on DA decay tau. Statistical analysis included all comparisons, but data are separated for clarity. DA decay was significantly higher in male than female WT mice, but there were no differences from, and no sex differences within LKI mice. There were also no significant sex differences between VKI mice, however trends underlie significant differences in DA decay of both male and female VKI, relative to WT, in the opposite direction of basal sex differences. LRRK2 inhibition had no effect on LKI or VKI DA decay (not shown) but significantly decreased and increased male & female WT PPRs, respectively, reversing base sex differences.

We first assessed glutamate release in response to increasing stimulation intensities. A significant interaction between genotype and stimulation intensity was observed, with reduced glutamate response peak in LKI and VKI mice, relative to WT controls (Figure 3Bi). *Post hoc* comparisons revealed significant reductions at higher stimulation intensities in LKI mice. No sex differences in peak glutamate responses were observed (not shown). To further assess presynaptic release properties, we examined glutamate paired-pulse ratios (PPRs; 100ms IPI), widely believed to indicate initial presynaptic probability of release (Pr)^18,19^. Although PPRs did not significantly differ between genotype, WT and LKI females exhibited significantly lower PPRs than their male counterparts, a sex difference absent in VKI (Figure 3Bii). Following MLi-2 treatment, glutamate PPRs were significantly reduced in WT males but not females, eliminating the baseline sex difference. A similar pattern was observed in LKI mice, but not VKI mice.

We next examined evoked DA release in aged WT, LKI, and VKI mice. A highly significant main effect of genotype was observed, with reduced DA responses across stimulation intensities in both LKI and VKI mice, relative to WT controls (Figure 3Ci). No significant sex differences in peak DA responses were detected (not shown). ANOVA revealed a highly significant effect on maximal DA response peaks, and *post hoc* multiple comparison testing found significant reductions in LKI males and VKI females relative to WT controls, with very strong trends toward reduced release in LKI females and VKI males (Figure 3Cii). No significant differences in maximal DA release were observed between vehicle-treated males and females within any genotype. LRRK2 inhibition significantly reduced maximal DA responses in WT females, but not males, and there were no significant treatment effects observed in LKI or VKI mice.

WT females exhibited significantly higher PPRs than WT males, whereas again, no significant sex differences or genotype-dependent alterations were observed in LKI or VKI mice (Figure 3Ciii). LRRK2 inhibition significantly increased PPRs in WT males, and decreased PPRs in WT females, reversing the baseline sex difference. In contrast, MLi-2 had no significant effect on DA PPRs in either LKI or VKI mice (not shown).

DA transient decay kinetics (tau) showed WT males had significantly slower responses than WT females, whereas but again no significant sex differences were observed in LKI or VKI mice (Figure 3Civ). Although decay kinetics did not significantly differ between WT and LKI mice, both male and female VKI mice exhibited significant differences relative to their respective WT controls, in directions opposite to the basal sex differences in WT mice. LRRK2 inhibition also produced opposing effects on DA decay kinetics in WT males and females, reducing decay tau in males and increasing it in females, thereby reversing the baseline sex difference. No significant treatment effects were observed in LKI or VKI mice (not shown).

Overall, the results demonstrate that both LKI and VKI mice exhibit altered striatal glutamate and DA neurotransmission at 25 months of age. Glutamate impairments were clearer in LKI mice than VKI, but subtle, and sensitive to LRRK2 inhibition. DA release was clearly impaired (30-50%) across a range of stimulus intensities in LKI & VKI, and similarly reduced in male and female VKI & LKI mice. Although LRRK2 inhibition consistently altered several measures of neurotransmission in aged WT mice, in a sex dependent manner, it did not in LKI or VKI mice, and it failed to reverse impaired DA release in aged LKI & VKI mice.

### Acute LRRK2 kinase inhibition restores dopamine release in 6-month-old LKI mice

To determine if deficits in striatal DA release can be reversed by LRRK2 inhibition, we examined the effect of MLi-2 in 6-month-old WT and LKI mice, the age at which we first detected impairments. Following acute brain slice preparation, paired slices from each animal were incubated in either vehicle (Captisol®) or MLi-2 (500nM) for 2h, then maintained under their respective treatment conditions during recording (Figure 4A). This paired experimental design enabled direct comparisons of DA release between vehicle- and MLi-2-treated slices from the same animal.

**Figure 4.**
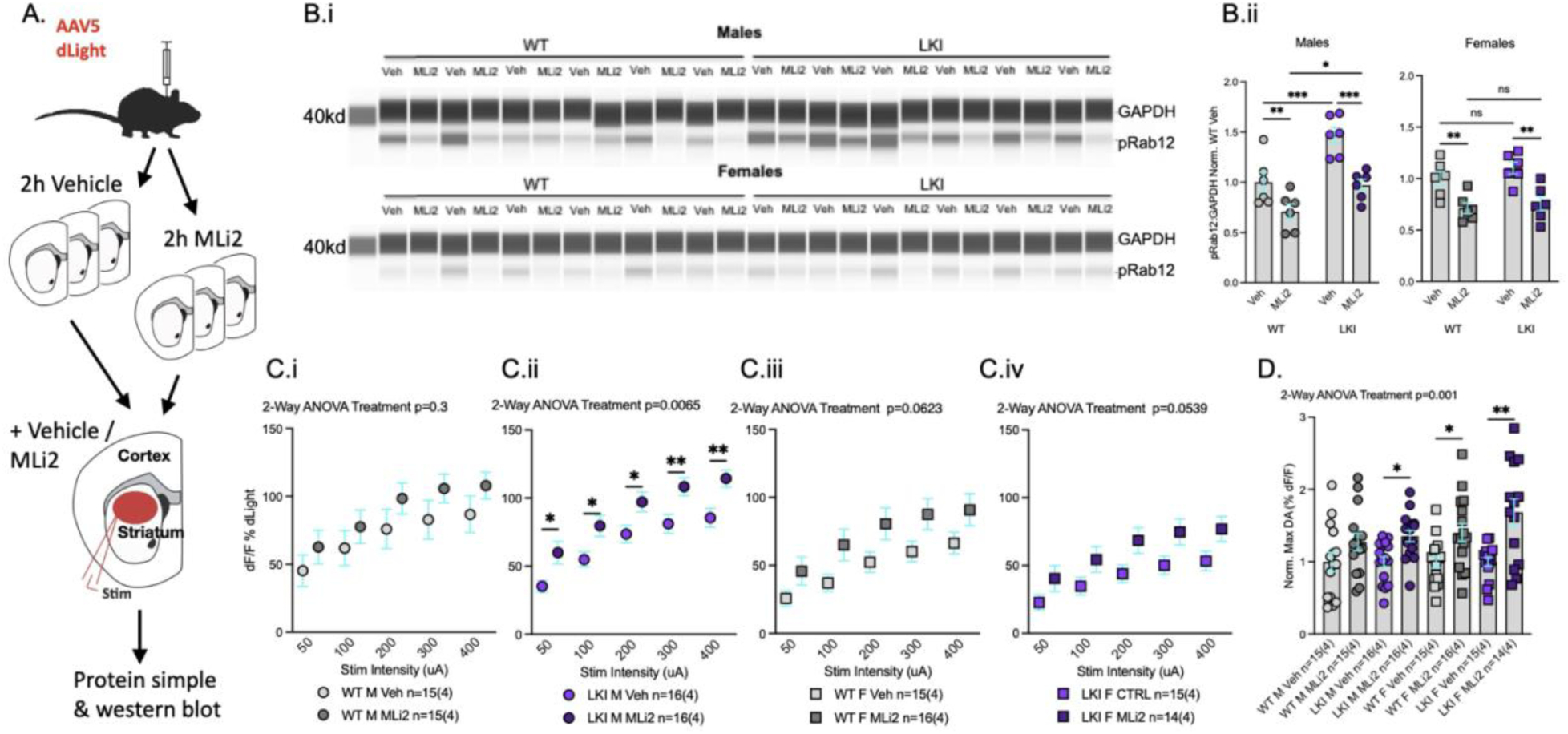
Dopamine release deficits at 6 months are rapidly reversed by acute LRRK2 inhibition. **A)** Schematic depicting stereotaxic injection of AAV5-dLight to dorsolateral striatum (DLS) 3–6 weeks prior to acute slice preparation, interleaved, paired slices from each animal are pre-incubated with vehicle (Captisol®) or MLi-2 (5500nM MLi-2) for 2h after brain slicing, and maintained in vehicle or MLi-2 during recording, prior to harvesting for biochemical assays. **B.i)** Raw data from Wes capillary-based protein separation and probe against GAPDH and pRab12 in slices from male (top) and female (bottom) WT & LKI mice. **ii)** Quantification (normalized to mean value of WT vehicle) shows effective LRRK2 inhibition by reduced pRab12. **Ci-iv)** There was no significant difference in raw DA release plots between vehicle and MLi-2 treated male (i) or female (iii) WT, nor LKI females (i); conversely DA release was significantly increased at all stimulation intensities in MLi-2 treated LKI males (ii). **D)** The experimental design enables pairing of MLi-2 treated and untreated slices from each animal. Within animal normalization of DA release maxima confirmed MLi-2 induced a significant increase of DA release in LKI males and revealed significant increases to DA release in WT & LKI females induced by LRRK2 inhibition.

To confirm effective LRRK2 inhibition, we assessed phosphorylation of the downstream LRRK2 substrate Rab12 (S106) using automated capillary based western blot on striatal slices collected following recording (Figure 4Bi). Quantification of pRab12 levels, normalized to the mean of WT vehicle controls, showed significantly reduced pRab12 with MLi-2 treatment, confirming target engagement and block of LRRK2 kinase activity (Figure 4Bii).

Prior to examining protein levels in slices, we evaluated LRRK2 inhibition and DA release. In WT mice, MLi-2 did not significantly alter DA responses in either males or females when comparing raw DA responses across stimulation intensities, although low to strong trends were apparent, respectively (Figure 4Ci,iii). Similarly, inhibition increased DA release in LKI females (Figure 4Civ), sitting on the border of statistical significance. In contrast, MLi-2 significantly increased evoked DA release across all stimulation intensities in LKI males, relative to vehicle-treated controls (Figure 4Cii).

To account for variability in baseline DA release between animals, we normalized maximal DA responses to the corresponding vehicle-treated slices within each animal (Figure 4D). This confirmed a significant increase in maximal DA release following LRRK2 inhibition in LKI males. Additionally, within-animal normalization revealed LRRK2 inhibition induced a significant increase in maximal DA release in WT and LKI females. An increase observed in WT males was not so profound as to reach statistical significance.

Collectively, the results establish that acute LRRK2 kinase inhibition enhances evoked striatal DA release in 6-month-old WT mice, with the clearest effect in females. This suggests ongoing LRRK2 activity has a physiological role negatively regulating DA release in the adult striatum. Furthermore, LRRK2 inhibition significantly potentiated DA release in male & female LKI mice, effectively reversing the DA release deficit phenotype.

## Discussion

We investigated how pathogenic LRRK2 variants alter striatal neurotransmission across ageing, and whether the functional changes are influenced by sex and ongoing LRRK2 kinase activity. Our data demonstrate that LRRK2-G2019S produces early, sex-dependent impairments in presynaptic DA signaling, with male LKI mice exhibiting pronounced deficits by 6 months of age. These alterations were reversible by acute LRRK2 kinase inhibition, supporting a role for elevated LRRK2 kinase activity in early dopaminergic dysfunction in PD variant scenarios, and strengthening the case for the neuroprotective potential of LRRK2 inhibitors. In contrast, aged LKI and VKI mice exhibited significant reductions in glutamate and DA transmission, but the latter was impervious to LRRK2 inhibition. These observations highlight the importance of age and sex when trying to understand the role of LRRK2 activity and PD variant effects on striatal neurotransmission, and when to attempt LRRK2 inhibition.

Interestingly, kinetic data suggest less-synchronous release and/or impaired DA clearance precedes reductions in DA release peaks. The dominant determinant of DA decay kinetics is clearance of extracellular DA by reuptake through DAT^25^; however, temporally high-resolution recording techniques show that (although most DA release events produce a single classical spike-like transient) 15–20% of release events consist of a succession of spikes that gradually decrease in amplitude ^25,26^. These “complex” events are believed to be due to flickering of the vesicle fusion pore, with each “flicker” releasing 25–30% of vesicular DA. A change in complex release would logically also alter DA kinetics. At 3 months of age, male LKI mice exhibited slower DA decay kinetics than WT males, but no reduction of DA release amplitude. By 6 months, this difference in decay was no longer observed in males, coinciding with the emergence of pronounced deficits in DA release. In contrast, female LKI mice exhibited no differences in decay kinetics at 3 months, but showed significantly slower decay than WT females and LKI males at 6 months, when reductions in DA release were less pronounced than in males. These insights suggest that increased decay times emerge in a sex- and age-dependent manner, and precede more substantial impairments in vesicular DA release. This raises the possibility that dysregulation of DA reuptake is an early component of LKI synaptic dysfunction; however, this is not supported by our previous report of increased decay tau in 3 month old LKI males, observed by voltammetry, which was conserved even in the presence of DAT block ^19^. We posit that LRRK2 hyperactivity (which is always present in the mice), initially slows responses by changing release properties, perhaps with more complex events, and that this (over time) transitions to more general DA release impairments.

Sex-dependent differences we observed in 6-month-old LKI mice, where DA release was significantly reduced in males, but relatively preserved in females, are consistent with previous reports of sex differences in presynaptic DA regulation. Females rely more heavily on VMAT2 vesicular repackaging to sustain DA release, and exhibit tighter regulation of extracellular DA by elevated DAT-mediated clearance ^27–29^. This may confer resilience against early impairments in DA neurotransmission caused by the G2019S variant.

A potential mechanism underlying impaired vesicular release is disruption of presynaptic membrane trafficking by LRRK2-G2019S. LRRK2 has been implicated in clathrin-mediated endocytosis (CME) and activity-dependent synaptic vesicle recycling, processes required to maintain the pool of release-competent vesicles during neurotransmission ^30,31^. Consistent with this, pathogenic LRRK2 variants have been associated with disrupted CME and synaptic defects in DA neurons, while activity-dependent endocytosis is reduced in iPSC-derived DA neurons carrying LRRK2 variants ^30^. Disruption of these pathways could therefore contribute to the reduced evoked DA release observed in LKI mice by impairing vesicle retrieval and recycling at the presynaptic terminal.

To further investigate the molecular mechanisms underlying these alterations in presynaptic DA regulation, we examined Rab12, an activator, and downstream substrate of LRRK2-dependent phosphorylation. Hyperphosphorylation of Rab GTPases is a well-established consequence of pathogenic LRRK2 & VPS35 activity in PD, although the downstream mechanisms contributing to neuronal dysfunction remain incompletely understood. Previous studies have demonstrated that Rab12 phosphorylation promotes LRRK2 recruitment to cellular membranes, facilitating the phosphorylation of additional Rab substrates and potentially amplifying LRRK2-dependent signaling ^32–34^. More recently, pRab12 has been shown to be enriched in synaptosomes and associated with docked synaptic vesicles at the presynaptic active zone, suggesting a potential role in the regulation of synaptic transmission ^35^. Consistent with this possibility, Rab12 overexpression has been shown to inhibit synaptic vesicle exocytosis, although whether Rab12 hyperphosphorylation produces similar effects remains unclear ^35^. These findings raise the possibility that dysregulated LRRK2–Rab12 signaling contributes to early synaptic vulnerability by interfering with presynaptic vesicle release mechanisms. This interpretation is consistent with our observation of reduced evoked DA release in LKI & aged VKI mice.

Whilst the results suggest a role for dysregulated LRRK2 signaling in presynaptic DA dysfunction, the effects of pathogenic LRRK2 variants also extend beyond dopaminergic terminals to disrupt glutamatergic neurotransmission ^12,19,36^. Our data demonstrate that glutamatergic responses are reduced in aged LKI and VKI mice, consistent with our previous work showing initially increased and age-dependent decline in glutamate transmission in LKI mice^19^. One potential mechanism underlying these deficits is disrupted trafficking of glutamate transporters and receptors. LRRK2-G2019S has been shown to impair trafficking of the excitatory amino acid transporter 2 (EAAT2), an effect reversed by MLi-2 treatment ^37^.

The present study also revealed sex-dependent differences in presynaptic glutamate release, with reduced paired-pulse ratios (PPRs) in WT and LKI females, compared with males. MLi-2 eliminated these differences by reducing PPRs in males, suggesting that LRRK2 kinase activity may influence presynaptic glutamate release in a sex-dependent manner. We previously observed reduced PPRs in striatal spiny projection neurons (SPNs) in 6-month-old VKI male mice ^18^. However, MLi-2 did not alter PPRs in WT or VKI SPNs^18^, nor alterations to spontaneous excitatory postsynaptic currents in cultured VKI neurons^10^. This suggests LRRK2 kinase inhibition differentially affects glutamatergic transmission in aged WT mice than in younger animals^18^, and has little effect on glutamatergic abnormalities in VKI. Taken together, these observations indicate that LRRK2 variants may impair glutamatergic signaling through distinct mechanisms involving glutamate homeostasis, postsynaptic receptor trafficking, and presynaptic release regulation.

In contrast to the therapeutic responsiveness observed at earlier ages, LRRK2 kinase inhibition failed to restore DA release in aged LKI and VKI mice. At 25 months, both genotypes exhibited significantly reduced evoked DA release across all stimulation intensities relative to WT controls. Maximal DA responses were significantly reduced in LKI males and VKI females, with strong trends to reductions in LKI females and VKI males. Age-dependent reductions in DA transmission in LKI mice have been previously reported, although findings differ regarding the nature of reductions, and age at which these deficits emerge. Previous studies have reported no differences in evoked extracellular DA between WT and LRRK2-G2019S overexpressing mice at 6 months of age, but reduced DA transmission by 12 months ^38^, and we ourselves showed evidence of early increases in repeated DA release that were absent at 12 months by voltammetry ^19^, and reduced by more sensitive dLight recording at 6 months here.

We have shown in three separate studies that VKI mice exhibit increased DA release at young 3-6 month ages ^18,21,23^. This contrasts with decreased DA release in LKI at 6 months and suggests, despite similarities in LRRK2 hyperphosphorylation, that LRRK2 G2019S & VPS35 D620N mechanisms disrupting DA function are related, but distinct. The G2019S mutation increases all LRRK2 activity, hyperphosphorylating many Rab GTPases crucial to regulation of the endolysosome system, synaptic vesicles, and synaptic function^39^. In contrast, the D620N variant in VPS35 is thought to secondarily activate LRRK2 through increased endolysosomal membrane stress^40^, which then activates a specific (if potentially deleterious) LRRK2 response. Further studies comparing the phosphoproteome in both models, at various ages, and in the presence of LRRK2 inhibition, may untangle common from distinct mechanisms. Regardless, LRRK2 inhibition reversed decreased release in LKI & increased release in VKI at early time points, arguing for a benefit in both scenarios.

Despite pronounced deficits observed in aged LKI and VKI mice, MLi-2 treatment did not improve DA release in either genotype, although it significantly reduced maximal DA responses in WT females. Similarly, LRRK2 inhibition did not alter DA PPRs or decay kinetics in aged LKI or VKI mice, despite producing sex-dependent changes in both measures in WT animals. These results suggest that LRRK2-associated dopaminergic dysfunction becomes less responsive to acute kinase inhibition with advancing age, potentially reflecting the accumulation of downstream alterations that are no longer readily reversible through suppression of ongoing LRRK2 kinase activity. Importantly, this contrasts with restorative effects observed at 6 months, arguing that ongoing LRRK2 activity is key to DA dysfunction in adult animals, and that it can be successfully mitigated if conducted in early stages. Regardless of no effective reversal of release deficits in aged LKI & VKI, chronic LRRK2 inhibition may still have neuroprotective benefits, and prevent additional progressive functional decline. The data support the therapeutic potential of targeting LRRK2 activity during prodromal and early stages of PD.

Nevertheless, a major challenge for clinical translation remains the reliable identification of prodromal PD. Consequently, the development of early diagnostic approaches remains an important area of research, with LRRK2-associated biomarkers currently under investigation. Overall, this work supports the growing body of evidence that dopamine axon dysfunction precedes retrograde and overt neurodegeneration in PD. By identifying early, sex-dependent alterations in presynaptic DA release, this study advances understanding of how LRRK2 kinase hyperactivity contributes to synaptic dysfunction and demonstrates the deficits are (at least partially) rapidly reversible by acute kinase inhibition. Together, the results expand our understanding of LRRK2 & VPS35 fundamental neurobiology, highlight the importance of sex and age in preclinical studies, and guide neuroprotective strategies aimed at preserving dopaminergic function before irreversible degeneration.

